# Widespread emergence of OmpK36 loop 3 insertions among multidrug-resistant clones of *Klebsiella pneumoniae*

**DOI:** 10.1101/2022.02.07.479342

**Authors:** Sophia David, Joshua L C Wong, Julia Sanchez-Garrido, Hok-Sau Kwong, Wen Wen Low, Fabio Morecchiato, Tommaso Giani, Gian Maria Rossolini, Stephen J Brett, Abigail Clements, Konstantinos Beis, David Aanensen, Gad Frankel

## Abstract

Mutations in outer membrane porins act in synergy with carbapenemase enzymes to increase carbapenem resistance in the important nosocomial pathogen, *Klebsiella pneumoniae* (KP). A key example is a di-amino acid insertion, Glycine-Aspartate (GD), in the extracellular loop 3 (L3) region of OmpK36 which constricts the pore and restricts entry of carbapenems into the bacterial cell. Here we combined genomic and experimental approaches to characterise the diversity, spread and impact of different L3 insertion types in OmpK36. We identified L3 insertions in 3588 (24.1%) of 14,888 KP genomes with an intact *ompK36* gene from a global collection. GD insertions were most common, with a high concentration in the ST258/512 clone that has spread widely in Europe and the Americas. Aspartate (D) and Threonine-Aspartate (TD) insertions were prevalent in genomes from Asia, due in part to acquisitions by ST16 and ST231 and subsequent clonal expansions. By solving the crystal structures of novel OmpK36 variants, we found that the TD insertion causes a pore constriction of 41%, significantly greater than that achieved by GD (10%) or D (8%), resulting in the highest levels of resistance to selected antibiotics. In a murine pneumonia model, KP mutants harbouring L3 insertions have a competitive disadvantage relative to a strain expressing wild-type OmpK36 in the absence of antibiotics. This explains the reversion of GD and TD insertions observed at low frequency among KP genomes. Finally, we demonstrate that strains expressing L3 insertions remain susceptible to drugs targeting carbapenemase-producing KP, including novel beta lactam-beta lactamase inhibitor combinations. This study provides a contemporary global view of OmpK36-mediated resistance mechanisms in KP, integrating surveillance and experimental data to guide treatment and drug development strategies.

**Author summary:** Rapidly rising rates of antibiotic resistance among *Klebsiella pneumoniae* (KP) necessitate a comprehensive understanding of the diversity, spread and clinical impact of resistance mutations. In KP, mutations in outer membrane porins play an important role in mediating resistance to carbapenems, a key class of antibiotics. Here we show that resistance mutations in the extracellular loop 3 (L3) region of the OmpK36 porin are found at high prevalence among clinical genomes and we characterise their diversity and impact on resistance and virulence. They include amino acid insertions of Aspartate (D), Glycine-Aspartate (GD) and Threonine-Aspartate (TD), which act by decreasing the pore size and restricting entry of carbapenems into the bacterial cell. We show that these L3 insertions are associated with large clonal expansions of resistant lineages and impose only a low fitness cost. Critically, strains harbouring L3 insertions remain susceptible to novel drugs, including beta lactam-beta lactamase inhibitor combinations. This study highlights the importance of monitoring the emergence and spread of strains with OmpK36 L3 insertions for the control of resistant KP infections and provides crucial data for drug development and treatment strategies.

## Introduction

*Klebsiella pneumoniae* (KP) is a leading cause of opportunistic infections in hospital and healthcare-associated settings worldwide (1,2). Rates of antibiotic resistance among KP have risen rapidly in recent years, leading to its classification by the World Health Organisation as a critical priority resistant pathogen (3). Of particular concern is the increasing number of KP infections that are resistant to carbapenems, which have been shown to be associated with a high mortality burden (4). Whilst newer agents with activity against carbapenem-resistant KP have been recently licensed (5), carbapenems remain vital in the treatment of severe infections due to their broad efficacy and limited adverse effects.

Carbapenem resistance in KP is primarily achieved by the acquisition of carbapenemase enzymes, which inactivate carbapenems by hydrolysis. These enzymes are typically plasmid-encoded and include variants of the KPC, OXA-48-like, NDM, VIM and IMP families (6). Another important mechanism involves the modification of outer membrane porins which enable the non-selective diffusion of substrates, including both nutrients and antibiotics, into the bacterial cell (7,8). These modifications restrict antibiotic entry and act in synergy with carbapenemase enzymes to increase the level of resistance. Resistance-associated mutations have been described in both major chromosomally-encoded KP porins, OmpK35 and OmpK36 (9,10). In particular, truncations in the *ompK35* gene that result in a non-functional porin have been widely identified and are ubiquitous in a major high-risk clone comprising sequence types 258 and 512 (ST258/512) (11,12). By contrast, *ompK36* is rarely truncated and resistance mutations more commonly either reduce OmpK36 abundance (13–15) or constrict the pore size (12,16).

Mutations that mediate pore constriction have been shown to consist of amino acid insertions in extracellular loop 3 (L3) of OmpK36, a motif that conformationally determines the minimal pore radius. In particular, we previously determined the crystal structure of OmpK36 with a Glycine-Aspartate (GD) L3 insertion and showed that this results in a 10% reduction in minimal pore diameter. This led to a 16-fold increase in the minimum inhibitory concentration (MIC) of meropenem, the most widely used carbapenem (16). *In silico* structural modelling has also predicted OmpK36 pore constriction by other L3 insertions (Threonine-Aspartate (TD) and Serine-Aspartate (SD)) observed in clinical KP genomes (12). However, in the absence of solved structures, important derived metrics (e.g., minimal pore diameter) of these porin variants remain incomplete, precluding rational drug design. Moreover, studies assessing the prevalence of L3 insertions among clinical isolates have been restricted to the identification of *a priori* defined L3 insertions (i.e. GD/TD in Lam et al. 2021 (10)) and/or limited by the temporal and geographic breadth of available sample collections (10,12). The increased availability of genomes now facilitates a more comprehensive analysis of different L3 insertions found among clinical KP worldwide, providing a valuable opportunity for informing surveillance, treatment and drug development strategies.

Here we used a combination of bioinformatic and experimental approaches to detail the diversity, evolutionary dynamics and clinical impact of L3 insertions observed among KP isolates from a large global genome collection (*n*=16,086). By solving additional OmpK36 structures, we show that other major L3 insertions beyond GD constrict the pore size and increase carbapenem resistance, driving large clonal expansions among high-risk clones. These include Aspartate (D) and TD insertions that are largely associated with the important albeit less well-studied ST16 and ST231 lineages, respectively. We also demonstrate recurrent reversions of L3 insertions among clinical isolates, in line with the competitive disadvantage of L3 insertion mutants observed in our preclinical mouse pneumonia model in the absence of antibiotic therapy. Finally, we systematically evaluated the effect of D, GD and TD insertions on the susceptibility to novel antibiotic therapies, including key beta-lactam/beta-lactamase inhibitor combinations, and show that these agents maintain efficacy despite pore constriction.

## Results

### D, GD and TD are the most common L3 insertions among clinical KP genomes

We first investigated the prevalence of L3 insertions among a large collection of public KP genomes available in PathogenWatch (https://pathogen.watch/genomes/all?genusId=570&speciesId=573; **S1 Table**). This collection comprises 16,086 assembled genomes from 84 countries, belonging to a total of 1,177 STs (17). We unambiguously identified the *ompK36* gene in 94.4% (15,193/16,086) of the genomes; the gene was intact (i.e. not truncated) in 98.0% (14,888/15,193) of these. Among those with an intact *ompK36*, we found that 24.1% (3588/14,888) had one or more amino acids inserted into the L3 region (**Figure 1A**). A total of eight different L3 insertion types were observed, which comprised between one and three amino acids. 75.3% (2700/3588) of the L3 insertions observed were GD, while the remainder comprised TD (14.3%; 512/3588), D (7.8%; 281/3588), SD (2.0%; 73/3588), N (0.4%; 15/3588), TYD (0.1%; 4/3588), YGS (0.06%; 2/3588) and GG (0.03%; 1/3588) (**Figure 1B)**.

**Figure 1.**
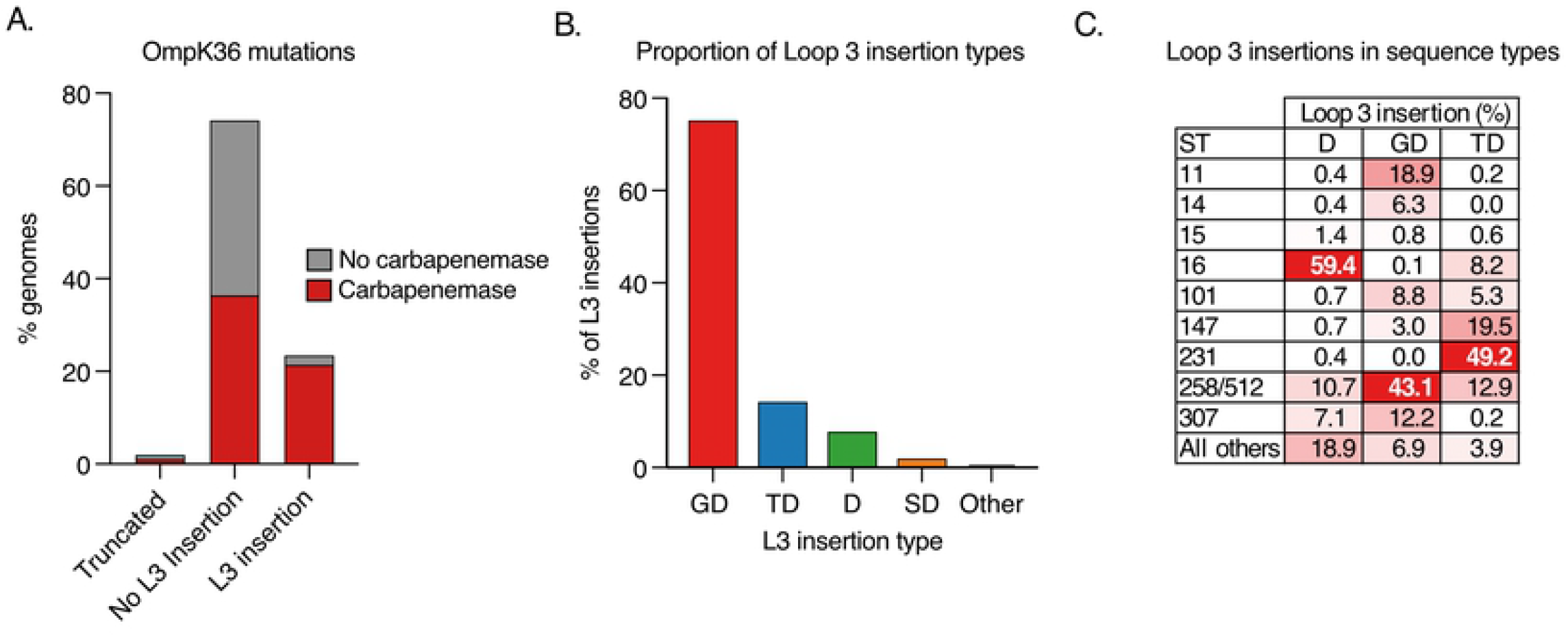
Diversity of loop 3 (L3) insertions among a collection of 16,086 public KP genomes. **A**. Proportion of genomes encoding an *ompK36* gene that is truncated or possessing a L3 insertion (if intact). Bars are coloured by the proportion of genomes carrying one or more carbapenemase genes. Only genomes with a single copy of *ompK36* that could be unambiguously characterised were included (*n*=15,193). **B**. Proportion of L3 insertions identified that comprise amino acid insertions of Glycine-Aspartate (GD), Threonine-Aspartate (TD), Aspartate (D), Serine-Aspartate (SD) or others (N/GG/TYD/YGS). **C**. Distribution of genomes possessing each L3 insertion type across major high-risk sequence types (ST). ST258 and ST512 are grouped together as they form a single clone.

We found L3 insertions in a total of 68 STs (GD - 52 STs, TD - 16 STs, D - 15 STs, SD - 10 STs, N - 1 ST, TYD - 3 STs, YGS - 2 STs), demonstrating their widespread emergence across the KP population. Notably, we found that the coding mutations for each L3 insertion were always the same, despite genetic redundancy (e.g. GD always encoded by ggc gac, TD by acc gac and D by gac). While this may partially be explained by exchange of alleles by recombination, phylogenetic analysis of the *ompK36* open reading frame (ORF) demonstrated parallel emergences of each insertion type across different gene backgrounds (**S1 Figure**). This suggests that these underlying coding mutations are more likely to evolve than the possible alternatives.

We found that L3 insertions were mostly concentrated among global multi-drug resistant (MDR) clones (**Figure 1C**), implicating an important role in resistance. Altogether, 91.8% (3295/3588) were found in one of the top ten most frequently observed STs in the genome collection, which represent these major clones. This is despite genomes from these STs making up only 56.2% (9045/16,086) of the total collection. We also observed a high concentration of individual L3 insertions in particular clones. For example, 43.1% (1163/2700) of genomes with a GD insertion belonged to either ST258 or ST512 (which together make up the single clone, ST258/512), 49.2% (252/512) of genomes with TD belonged to ST231, and 59.4% (167/281) of genomes with D belonged to ST16 (**Figure 1C**). While all three clones are internationally dispersed, ST258/512 has largely been a dominant strain in the Americas, Europe and Middle East (18–22), and ST231 and ST16 are found at high prevalence in parts of Asia (23–25).

We also found that L3 insertions frequently co-occur with carbapenemase genes, which have also been shown to be concentrated among major high-risk clones (26). Indeed, of the genomes in this collection with an L3 insertion, 90.7% (3255/3588) possessed one or more carbapenemases (**Figure 1A**). Furthermore, we observed that particular L3 insertions more frequently co-occur with some carbapenemases. For example, 76.0% (1854/2441) of carbapenemases found among genomes with a GD insertion were KPC, while 72.3% (334/462) and 75.7% (203/268) of those found among genomes with D and TD were from the OXA-48-like families, respectively. However, these associations are also confounded by the high concentration of particular carbapenemase genes among some lineages (e.g. KPC in ST258/512).

### The L3 insertions reduce meropenem diffusion while enabling glucose entry

We next aimed to define the extent to which the D and TD insertions, observed at highest prevalence after GD insertions, constrict the OmpK36 pore and increase meropenem resistance. To that end we solved the crystal structures of chimeric OmpK36_WT_ with D and TD insertions (OmpK36_WT+D_ and OmpK36_WT+TD_) by X-ray crystallography and compared the minimal pore diameters to those of the previously solved OmpK36_WT_ and OmpK36_WT+GD_ structures (**Figure 2A-D; S2 Table**). While the OmpK36_WT+D_ structure demonstrated the presence of two different L3 conformations, open or closed, both yielded similar minimal pore diameters (2.94 Å (D-open) and 2.95 Å (D-closed)). Notably, OmpK36_WT+TD_ forms a particularly narrow channel with a minimal pore diameter of 1.88 Å. These values represent relative pore reductions of 8% (D) and 41% (TD) compared to OmpK36_WT_, the latter significantly greater than that imposed by the GD insertion (10%).

**Figure 2.**
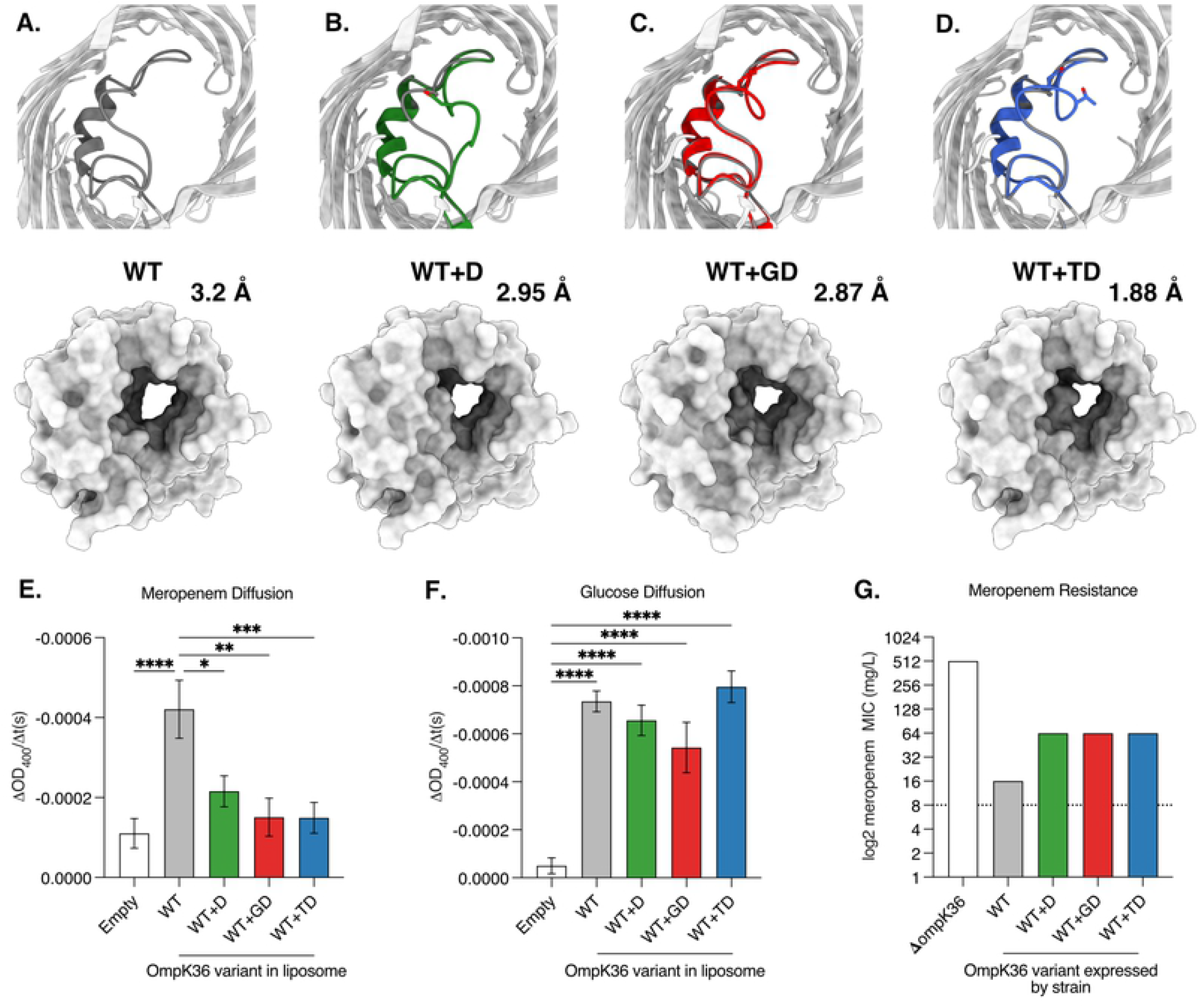
L3 insertions reduce OmpK36 pore diameter and restrict the diffusion of meropenem. **A-D.** Cartoon illustrations of OmpK36 (grey) in which L3 is coloured for all the variants; the view is from the extracellular space and perpendicular to the membrane (top panels). Insertions in L3 reduce the relative pore radius as calculated by the program HOLE (51); surface representation to show the impact of the mutations in the pore size (bottom panels). **E-F.** Rate of diffusion (ΔOD_400_/t(s) of meropenem (E) and glucose (F) as determined by liposome swelling assays for different OmpK36 variants. **G.** Median meropenem MIC values for strains encoding different *ompK36* variants (*n*=3 replicates). **E-F**: error bars ±SEM. * p<0.332; ** p<0.0021, *** p<0.0002, **** p<0.00001, statistical significance determined by ordinary one-way ANOVA with Tukey’s multiple comparison test.

To evaluate the effect of the reduced pore diameter on meropenem diffusion in OmpK36_WT+D_ and OmpK36_WT+TD_, we conducted liposome swelling assays. We also included OmpK36_WT_ and OmpK36_WT+GD_ variants, as they had been previously validated in these assays, and empty liposomes were used as a control to establish the baseline diffusion that occurs in the absence of OmpK36 channels. Diffusion rates were calculated by assessing changes in OD_400nm_ per unit time (ΔOD_400_/t(s)). We found that liposomes with the OmpK36_WT+D_ and OmpK36_WT+TD_ variants had significantly reduced meropenem diffusion compared to OmpK36_WT_, with rates similar to those observed with OmpK36_WT+GD_ and empty liposomes (**Figure 2E**). We also measured the diffusion rate of glucose, a key carbon source, which has a substantially lower molecular weight than meropenem (180.2g/mol vs 383.5g/mol). We found similar glucose diffusion in the presence of all OmpK36 variants (**Figure 2F**), which was higher compared to empty liposomes.

Finally, to determine the effect of the different L3 insertions on meropenem MIC, we replaced the endogenous (genomic) *ompK36*_WT_ ORF in KP strain ICC8001 with alleles encoding D, GD and TD insertions. As a control, we generated a strain lacking *ompK36*, the deletion of which has been shown to increase carbapenem resistance (9). Given the high frequency of *ompK35* truncations among high-risk clones we deleted the *ompK35* gene from all isogenic strains, and introduced the KPC-2-encoding plasmid, pKpQIL, by conjugation (see **Table 1** for list of strains and attributes). All L3 insertions increased the meropenem MIC four-fold from 16mg/L in *ompK36*_WT_ to 64mg/L (the resistance breakpoint is >8mg/L). The absence of *ompK36* increased the MIC 32-fold to 512mg/L.

**Table 1.**
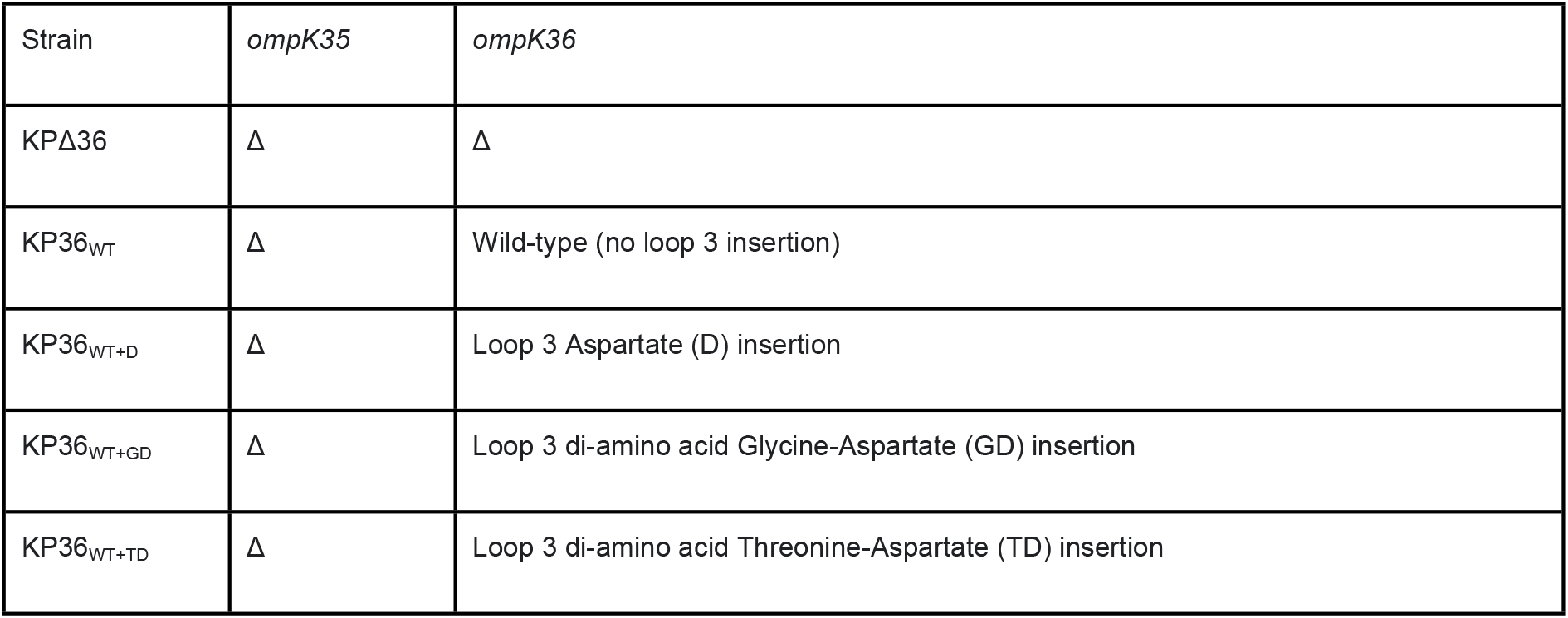
Isogenic strains used in this study. The parental strain ICC8001 (16) was genetically modified in a seamless and markerless recombineering approach to generate the *ompK36* variants.

### L3 insertions are associated with clonal expansions in MDR lineages and revert at a low frequency

We next investigated the emergence and expansion patterns of L3 insertions among clinical genomes using a phylogenetic approach. We analysed the three MDR clones in which the D, GD and TD insertions predominate (D - ST16; GD - ST258/512; TD - ST231) (**Figure 1B**). We included genomes from the Pathogenwatch collection belonging to each of these STs, which represented 3629 ST258/512 isolates (34 countries; collected 2003-2020), 446 ST16 isolates (26 countries; 2004-2020) and 302 ST231 isolates (19 countries; 2003-2019).

Our phylogenetic analysis of ST258/512 showed that an *ompK35* truncation and the KPC gene were largely ubiquitous and present in the earliest sampled isolates, while the lineage initially expanded in the absence of L3 insertions (**Figure 3A**). Since the emergence of this clone, L3 insertions have evolved many times independently. We found a total of six different L3 insertion types (D, N, GD, TD, SD, TYD) with D, GD and TD each emerging on multiple occasions. Several of the L3 insertion acquisition events (most notably of GD) are associated with subsequent clonal expansions. For example, the clade consisting largely of ST512 that encodes a GD insertion became highly successful as evident in the phylogeny and supported by multiple surveillance reports (18,27). Our data also confirmed previous reports that this clade likely spread from the USA to Europe and the Middle East (26) where it dominated the resistant KP population in some countries over several years (e.g. Italy and Israel) (18,27). Notably, the ST258/512 phylogeny also suggested that there have been multiple reversion events of L3 insertions, represented by genomes lacking a particular L3 insertion amidst a clade carrying that insertion (**Figure 3B**). The occurrence of reversions is suggestive of a selective pressure acting in favour of removing L3 insertions in certain contexts.

**Figure 3.**
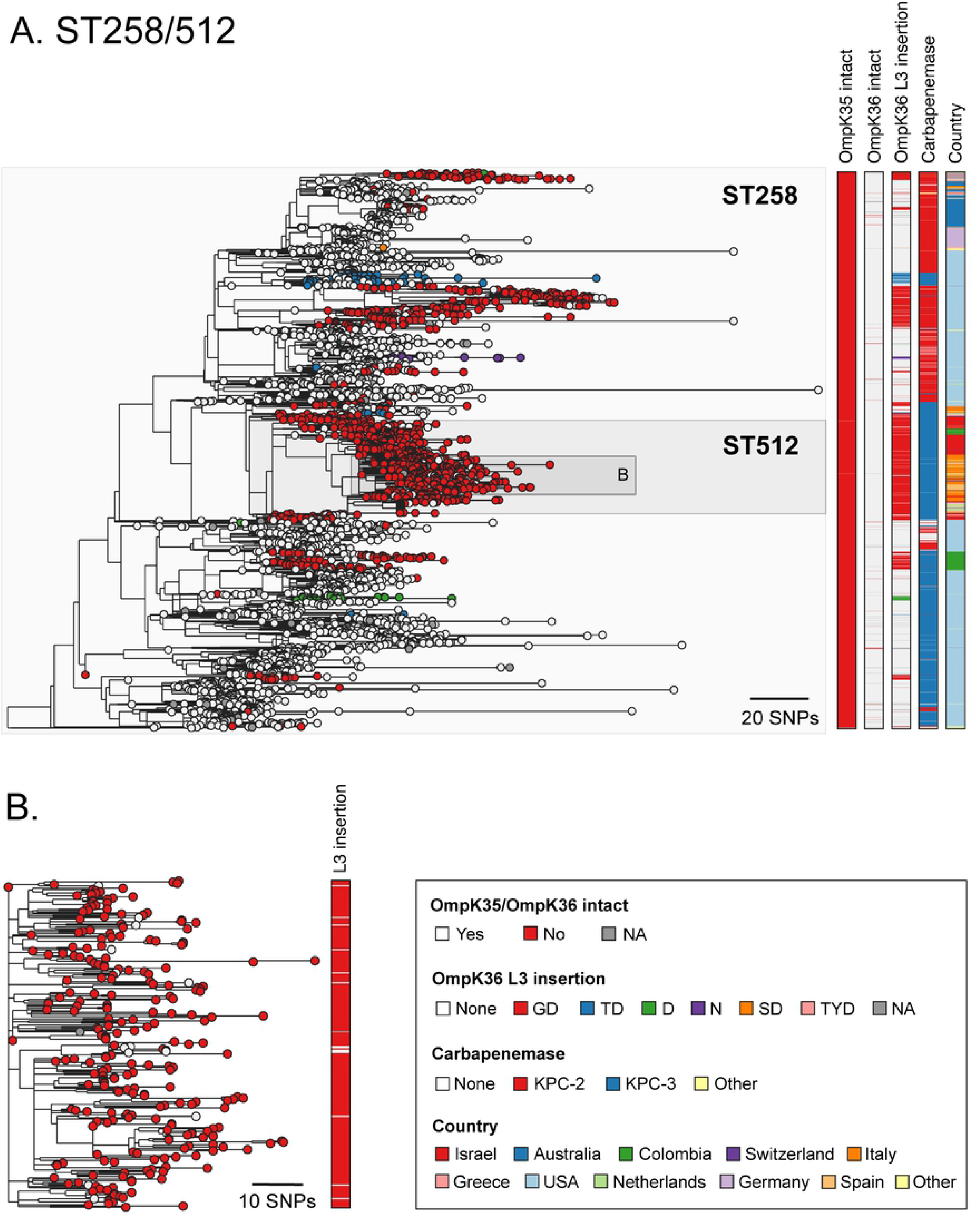
Multiple acquisitions of GD and other L3 insertions among the high-risk ST258/512 clone. **A.** Phylogenetic tree of 3629 isolates with public genome data belonging to sequence types (ST) 258 and 512. The tree was rooted using an ST895 isolate (accession SRR5385992) that was subsequently removed. Isolate tips are coloured by the type of OmpK36 L3 insertion (if applicable). Metadata columns from left to right show whether the *ompK35* and *ompK36* genes are intact (i.e., not truncated), the type of OmpK36 L3 insertion (if applicable), the carbapenemase gene type (if applicable) and the country of origin. Carbapenemases and countries are shown only for those with ≥15 isolates. Carbapenemase gene variants that have imperfect matches to known variants are grouped together with the most closely-related known variant. A similar interactive visualisation with more detailed metadata is available using Microreact at https://microreact.org/project/exB9brEAsQcpg7vKXMJtoF-k-pneumoniae-st258512 **B**. A zoomed-in visualisation of the clade highlighted in (A). Scale bars show the number of SNPs.

As with the ST258/512 lineage, our phylogenetic analysis of ST16 demonstrated that the lineage also expanded in the absence of L3 insertions and that these have since evolved frequently across different clades (**Figure 4A**). A total of five different L3 insertion types (D, GD, TD, SD, TYD) were found, with D, GD, TD and SD each evolving two or more times. We found a high diversity of carbapenemases among the ST16 lineage and numerous independent truncations of *ompK35*. However, most isolates (98.8%; 164/166) with the D insertion belonged to a single clade of isolates collected in Thailand between 2016-2018. This acquisition of the D insertion coincided closely with the gain of OXA-232 and NDM-1 carbapenemase genes and an *ompK35* truncation, likely leading to the rapid clonal expansion of this clade.

**Figure 4.**
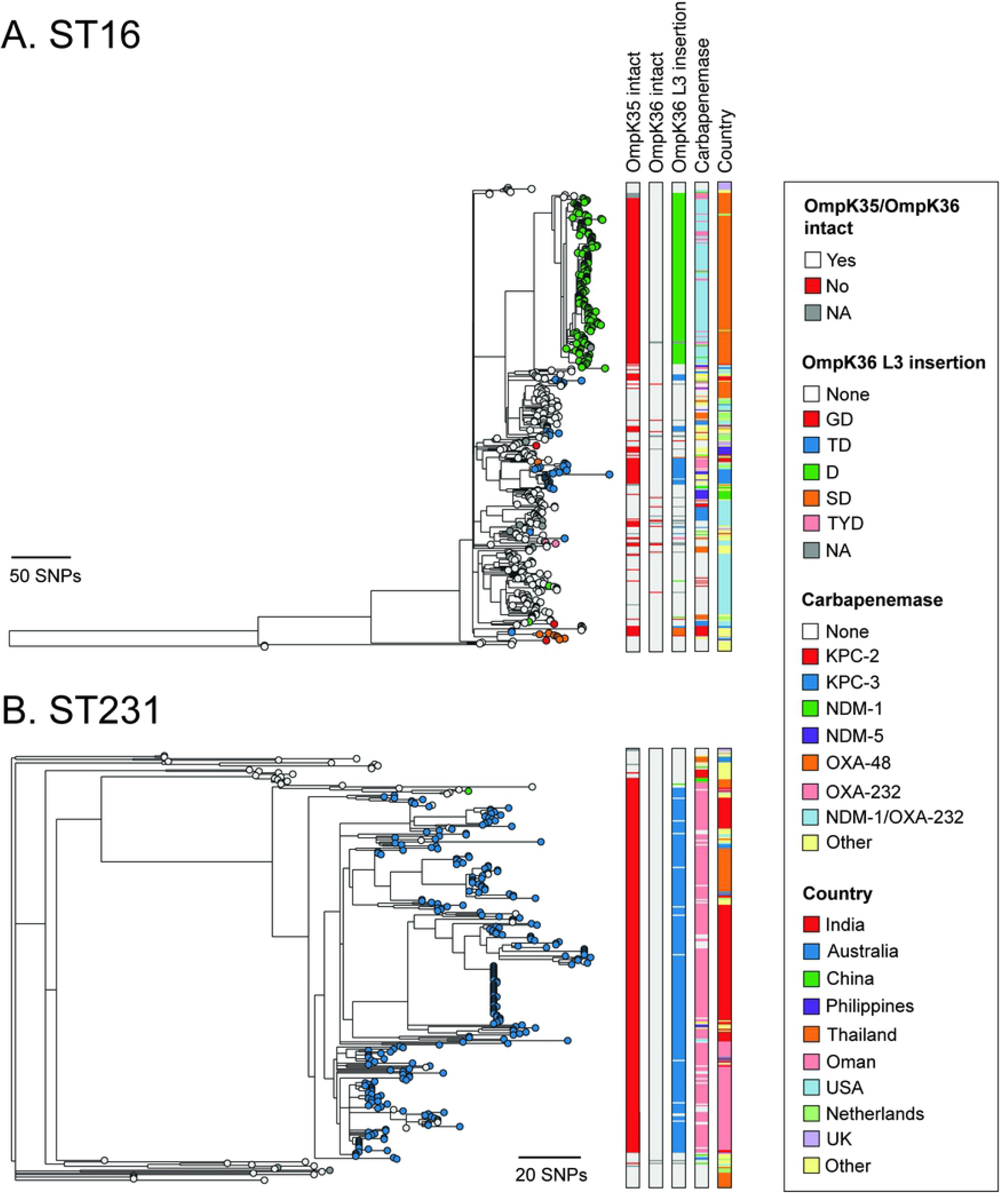
The D and TD insertions are associated with large clonal expansions in ST16 and ST231. Phylogenetic trees of 446 and 302 isolates with public genome data belonging to ST16 (**A**) and ST231 (**B**), respectively. The trees were rooted using an ST17 outgroup (accession ERR1228220) and an ST101 outgroup (accession ERR1216956), respectively, each of which were subsequently removed. Isolate tips are coloured by the type of OmpK36 L3 insertion (if applicable). Metadata columns from left to right show whether the *ompK35* and *ompK36* genes are intact (i.e., not truncated), the type of OmpK36 L3 insertion (if applicable), the carbapenemase gene type (if applicable) and the country of origin. Carbapenemases and countries are shown only for those with ≥10 isolates. Carbapenemase gene variants that have imperfect matches to known variants are grouped together with the most closely-related known variant. Scale bars show the number of SNPs. Similar interactive visualisations with more detailed metadata are available using Microreact at https://microreact.org/project/m8qd8j1YmfMapiPJ7prAEh-k-pneumoniae-st16 (ST16) and https://microreact.org/project/pZRm6DsxvZVYPQ2Ea33Buw-k-pneumoniae-st231 (ST231).

Contrary to the observations in ST258/512 and ST16, the ST231 phylogeny suggested that the TD insertion was acquired on a single occasion and associated with the major clade (**Figure 4B**). The low diversity within this clade is suggestive of a rapid clonal expansion. The acquisition of the TD insertion also coincided closely with the gain of OXA-232 and an *ompK35* truncation (with both likely occurring just prior to TD acquisition). No other L3 insertions were found in the ST231 lineage except for a single isolate with a D insertion. Phylogeographic analysis showed that the major clade encoding the TD insertion has spread to multiple countries, including India, Thailand and Oman, where significant local transmission is evident. As in the ST258/512 lineage, we also found numerous reversions of the TD insertion.

### The *in vivo* competitive disadvantage of L3 insertions explains the observed reversions

The high prevalence of the different OmpK36 L3 insertions across the KP population, together with our observation that reversions also occur, led us to explore the impact of the D, GD and TD insertions on bacterial fitness using a mouse pneumonia model. To do this, we performed *in vivo* infection experiments using isogenic strains of ICC8001 that encoded either *ompK36*_WT_ (KP36_WT_), *ompK36*_WT+D_ (KP36_WT+D_), *ompK36*_WT+GD_ (KP36_WT+GD_) or *ompK36*_WT+TD_ (KP36_WT+TD_) (**Table 1**). Intratracheal intubation was used to inoculate 250 CFU of KP directly into the lungs of mice, replicating ventilator-associated pneumonia (**Figure 5A**). A control group of animals received PBS alone. After 48 h, infection with all isogenic strains induced significant weight loss compared to those receiving PBS only (**Figure 5B**). However, no significant differences were observed between those infected with KP36_WT_ or KP36_WT+D_, KP36_WT+GD_ or KP36_WT+TD_. Similarly, all isogenic strains achieved high CFU counts in the lungs and blood (although not all animals were bacteraemic at the end of the time-course), with no significant differences observed between groups (**Figure 5C-D**). Measurement of proinflammatory cytokines revealed significant increases of serum IL-6 and CXCL-1 following infection with any strain compared to the PBS control, but with no significant differences observed between the strains themselves; however, serum TNF was only significantly elevated in KP36_WT_ infection compared to the uninfected controls (**Figure 5E-G**). Lastly, we found significant increases in lung neutrophils induced by all infecting strains compared to the PBS control, but observed no significant differences between the four OmpK36 backgrounds (**Figure 5H**). These experiments suggest that the L3 insertions do not significantly attenuate KP infection, thereby explaining the successful clonal expansions observed among isolates carrying these mutations.

**Figure 5.**
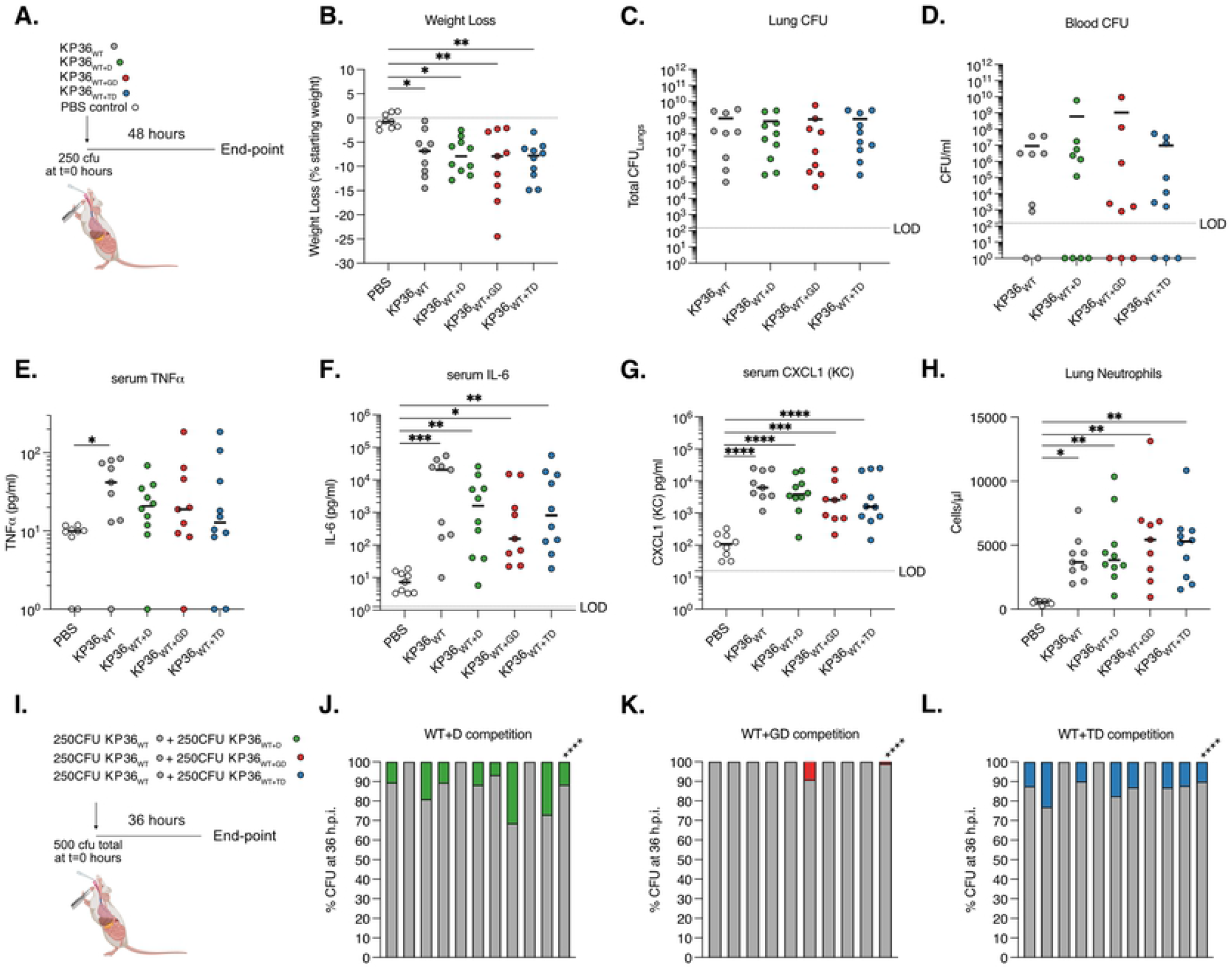
OmpK36 L3 insertions maintain virulence but confer a competitive disadvantage in a preclinical murine pneumonia model. **A-H.** Pneumonia was induced by the intratracheal administration of 250 CFU of isogenic KP strains expressing D (KP36_WT+D_), GD (KP36_WT+GD_) and TD (KP36_WT+TD_) OmpK36 L3 insertions. A strain lacking any L3 insertion (KP36_WT_) and PBS (uninfected) were used as controls. A schematic of the infection is outlined in panel **A**. At 48 hours post infection significant weight loss was induced by infection with all strains, irrespective of OmpK36 variant (**B)** and no significant differences were observed between strains. No significant differences were observed in the lung (**C)** and blood (**D**) bacterial burdens between infection with any strain. Serum TNF was only significantly increased following infection with KP36_WT_ compared to uninfected (PBS) controls (**E).** Serum IL-6 (**F**), CXCL-1 (**G**) and lung neutrophils (**H**) significantly increased following infection with all strains compared to uninfected (PBS) controls, with no significant differences between strains observed. **I-L.** Competition assays were employed to stringently assess the fitness of OmpK36 L3 insertion mutations. A schematic of the infection is outlined in panel **I.** 250 CFU of KP36_WT_ was competed against 250 CFU of KP36_WT+D_ (**J**), KP36_WT+GD_ (**K**) or KP36_WT+TD_ (**L**) and bacterial burdens were assessed at 36 hours post infection in the lungs. Each graph shows the % CFU recovered in the lungs in individual mice followed by a summary bar with the mean competition result across infections. KP36_WT_ significantly outcompeted all the L3 insertions tested. *, p < 0.05; **, p < 0.01; ***, p < 0.001; ****, p<0.0001. All experiments were conducted in biological duplicate with 4-5 mice per group. **B-H** Significance was determined by ordinary one-way ANOVA followed by Tukey’s multiple comparison post-test, except in **E** where Kruskal-Wallis test was employed as data was not normally distributed. **J-L** Mean competition was assessed by Mann-Whitney T-test.

We next used a more stringent method of assessing relative bacterial fitness by competing KP strains with L3 insertions against KP36_WT_ *in vivo*. We infected mice with a total inoculum of 500 CFU, comprising 50% KP36_WT_ and 50% of either KP36_WT+D_ or KP36_WT+TD_ (KP36_WT+GD_ having been tested previously (16) (**Figure 5I**). To identify the strains at the experimental end-point (36 hpi), we chromosomally tagged the input bacteria with either sfGFP or mRFP1. We enumerated lung CFU counts as the outcome measure and found that both KP36_WT+D_ and KP36_WT+TD_ with KP36_WT+GD_ previously) were outcompeted by KP36_WT_ (**Figure 5J-L**). These findings provide an experimental basis to explain the observed reversions of L3 insertions in the KP population as, whilst their expression supports a high capacity for infection, they result in a competitive disadvantage in the absence of antibiotics.

### Novel drugs targeting KPC-producing KP are effective against L3 insertion-expressing strains

Finally, we used our isogenic strain collection to systematically test the impact of D (KP36_WT+D_), GD (KP36_WT+GD_) and TD (KP36_WT+TD_) insertions on susceptibility to new or recently licensed antibiotic therapies that are vital for the treatment of carbapenemase-producing KP (**Table 2**). We evaluated four beta-lactam/beta-lactamase inhibitor combinations: ceftazidime/avibactam (CAZ/AVI), meropenem/vaborbactam (MER/VAB), imipenem/relebactam (IMI/REL) and aztreonam/avibactam (AZT/AVI)) and the novel siderophore cephalosporin cefedericol (FDC). When combination drugs were assessed, we also determined the MIC in the absence of the beta-lactamase inhibitor (i.e., parental drug alone). All strains had *ompK35* deleted and expressed the KPC-2 carbapenemase. We also evaluated the impact of deleting *ompK36* (KPΔ36) to replicate the effect of *ompK36* truncation, which was observed, albeit rarely, in our genomic analyses (**Figures 3A, 4A and 4B**).

**Table 2.**
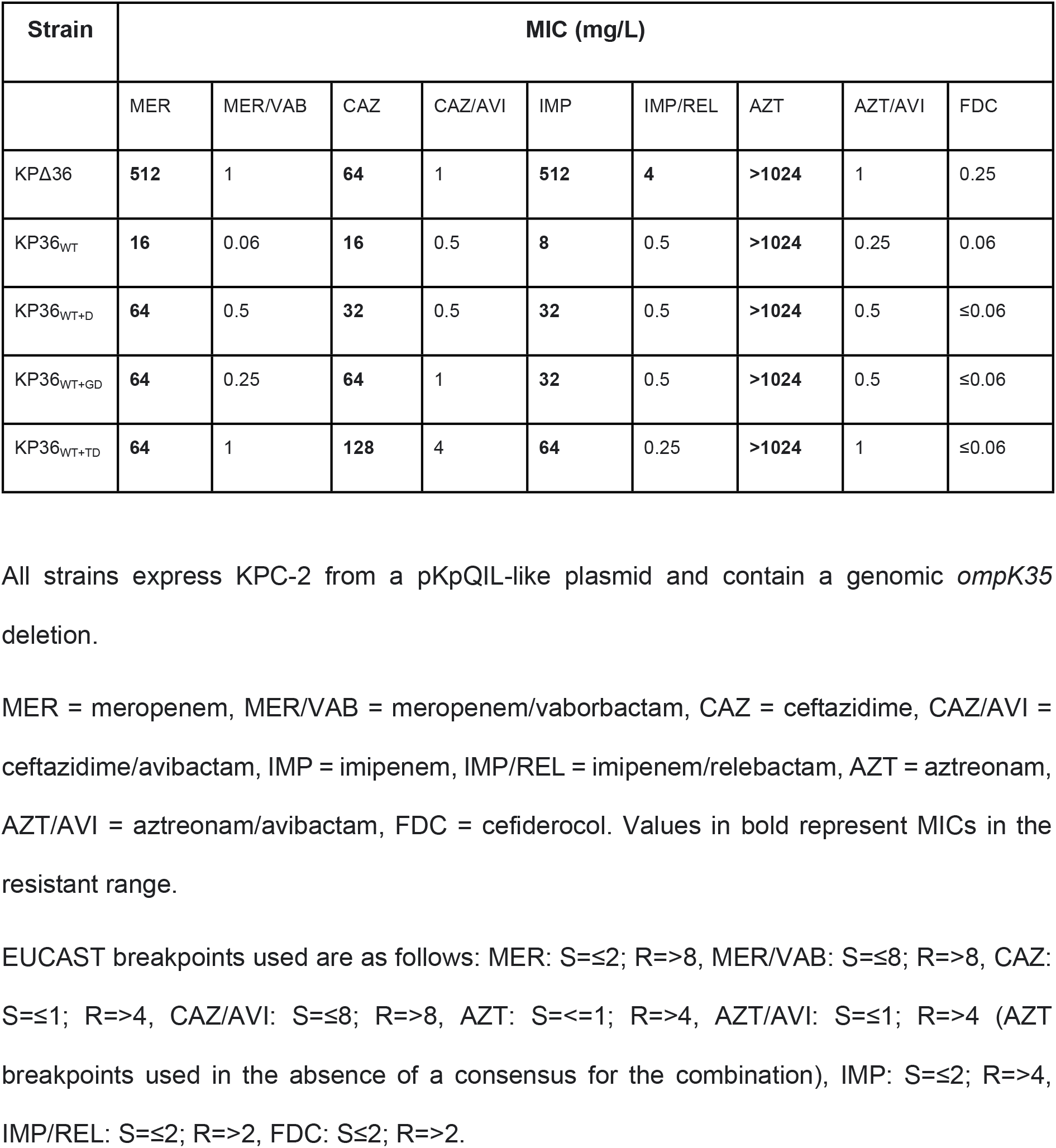
Minimum inhibitory concentrations of different antibiotics for isogenic KP strains.

| Strain | MIC (mg/L) |  |  |  |  |  |  |  |  |
| --- | --- | --- | --- | --- | --- | --- | --- | --- | --- |
|  | MER | MER/VAB | CAZ | CAZ/AVI | IMP | IMP/REL | AZT | AZT/AVI | FDC |
| KPΔ36 | **512** | 1 | **64** | 1 | **512** | **4** | **>1024** | 1 | 0.25 |
| KP36<sub>WT</sub> | **16** | 0.06 | **16** | 0.5 | **8** | 0.5 | **>1024** | 0.25 | 0.06 |
| KP36<sub>WT+D</sub> | **64** | 0.5 | **32** | 0.5 | **32** | 0.5 | **>1024** | 0.5 | ≤0.06 |
| KP36<sub>WT+GD</sub> | **64** | 0.25 | **64** | 1 | **32** | 0.5 | **>1024** | 0.5 | ≤0.06 |
| KP36<sub>WT+TD</sub> | **64** | 1 | **128** | 4 | **64** | 0.25 | **>1024** | 1 | ≤0.06 |

All strains express KPC-2 from a pKpQIL-like plasmid and contain a genomic *ompK35* deletion.
MER = meropenem, MER/VAB = meropenem/vaborbactam, CAZ = ceftazidime, CAZ/AVI = ceftazidime/avibactam, IMP = imipenem, IMP/REL = imipenem/relebactam, AZT = aztreonam, AZT/AVI = aztreonam/avibactam, FDC = cefiderocol. Values in bold represent MICs in the resistant range.
EUCAST breakpoints used are as follows: MER: S=≤2; R=>8, MER/VAB: S=≤8; R=>8, CAZ: S=≤1; R=>4, CAZ/AVI: S=≤8; R=>8, AZT: S=≤1; R=>4, AZT/AVI: S=≤1; R=>4 (AZT breakpoints used in the absence of a consensus for the combination), IMP: S=≤2; R=>4, IMP/REL: S=≤2; R=>2, FDC: S=≤2; R=>2.

The MICs to the parental drugs used in co-formulations (MER, CAZ, IMP and AZT) were in the resistant range, irrespective of the OmpK36 variant expressed (**Table 2**). As already described for MER (**Figure 2G**), D, GD and TD insertions increased the MIC as compared to wild-type OmpK36 expression. Whilst the MICs to AZT were universally above the range of the assay, CAZ and IMP resistance was found to be inversely correlated with the minimal pore diameter imposed by the L3 insertion type, with the KP36_WT+TD_ strain achieving higher MICs than seen in KP36_WT+D_ and KP36_WT+GD_. The KPΔ36 strain achieved the highest MICs for each drug with the exception of CAZ, where KP36_WT+TD_ obtained the highest level of resistance. When we tested these drugs in combination with their respective beta-lactamase inhibitors, susceptibility was restored among all strains harbouring L3 insertions. However, some increase was observed for the MICs of CAZ/AVI, MER/VAB and AZT/AVI (relative to KP36_WT_), suggesting that L3 insertions have some effect on the susceptibility to these novel beta-lactamase-inhibitor combinations. Of note, KPΔ36 presented a resistant phenotype to IMP/REL, which was not observed with MER/VAB, CAZ/AVI or AZT/AVI. All strains were susceptible to FDC, in keeping with a porin independent uptake mechanism via siderophore receptors (28). However, we noted that the FDC MIC was highest in KPΔ36 indicating that entry can, in part, be mediated by OmpK36. Taken together, these results have important medical implications as they show that novel drugs targeting carbapenemase-producing KP remain effective against strains possessing L3 insertions.

## Discussion

Monitoring the emergence, spread and clinical impact of resistance mutations among KP isolates is essential to informing public health intervention strategies. Here we demonstrate the widespread distribution of OmpK36 L3 insertions among clinical KP isolates worldwide, as inferred from a large collection of publicly available genomes (*n*=16,086). In particular, three types of L3 insertion, comprising amino acid insertions of D, GD and TD, made up 97.4% of those identified and were found in 23.5% (3493/14,888) of all genomes encoding an intact *ompK36* gene. Among genomes encoding one or more carbapenemase genes, this proportion increased to 36.1% (3171/8795). While there is a bias towards sequencing resistant isolates, our data nevertheless demonstrates that these mutations are one of the major adaptations of KP to antibiotic-rich healthcare environments.

We solved the structures of OmpK36 with the D and TD insertions and compared their minimal pore diameters to those of the wild-type OmpK36 porin and OmpK36 with a GD insertion determined previously (16). This revealed variation in the degree of pore constriction imposed by different L3 insertions, with reductions in pore size from the wild-type OmpK36 ranging from 41% (TD) to 10% (GD) to 8% (D). Interestingly, all three L3 insertions increased the meropenem MIC by the same magnitude (four-fold) compared to wild-type OmpK36 expression. However, the severity of pore constriction was reflected in the resulting MIC values for imipenem and ceftazidime (i.e. TD-expressing strain exhibiting the highest resistance). These differences are therefore vital in the process of rational physico-chemical drug design. We also found that L3 insertions have no effect on the diffusion of glucose, a key carbon source that is of lower molecular weight than beta-lactam antibiotics. The ability to maintain this physiological role of OmpK36 thereby demonstrates a key advantage of using pore constriction to impede antibiotic entry rather than mutations that reduce *ompK36* expression or result in a non-functional, truncated porin.

We showed that L3 insertions have emerged widely across the KP population and are most concentrated among known high-risk clones. In particular, ST258/512 accounted for a large proportion (43.1%) of GD insertions, while ST231 and ST16 harboured a high proportion of the TD (49.2%) and D (59.4%) insertions respectively. The relative lack of surveillance and availability of ST231 and ST16 genomes as compared to ST258/512 sequences can partially account for the under-recognition of the TD and D insertions to date, despite their high global clinical impact. Among these different clonal lineages, we found that L3 insertions often coincide with carbapenemases and *ompK35* truncations, and we found multiple instances of where the acquisition of these three traits in close succession was followed by rapid clonal expansion. Examples of this are the major clade of the ST231 lineage, associated with high transmission in India (and elsewhere) (23,25), as well as a large clonal expansion of an ST16 subtype in Thailand. We propose that these expansions may have been driven by the overuse of carbapenems prior to the availability of novel drugs specifically targeting carbapenemase-producing *Enterobacteriaceae* (CPE). In particular, high-dosage carbapenems in combination with other drugs (e. g. colistin, tigecycline, fosfomycin) or even the use of double carbapenem regimens were mostly recommended for treatment of CPE infections (29,30). Surveillance and infection control efforts must now focus on limiting spread of these resulting clones, as well as rapidly detecting the convergence of these resistance traits among other STs.

The association of L3 insertions with large clonal expansion events fits with our finding that these mutations do not reduce the infection capacity of KP in a mouse pneumonia model. This contrasts with mutations resulting in non-functional OmpK36, which cause even higher carbapenem resistance but have been shown to result in a large fitness cost (9,16) and rarely proliferate beyond individual patients. However, we did also find reversion events of L3 insertions (namely GD and TD) in clinical isolates, suggestive of a selection pressure acting to revert the pore to wild-type in certain environments (e.g., in the absence of antibiotics). This can be explained by the *in vivo* competition experiments performed here and previously (16), which demonstrated a competitive disadvantage of OmpK36 L3 insertion mutants relative to a wild-type strain. This suggests that antibiotic stewardship measures could play a crucial role in limiting further expansion of resistant KP carrying L3 insertions.

Finally, the high prevalence of L3 insertions among carbapenemase-producing strains led us to determine the precise impact of these mutations on the efficacy of new or recently licensed drugs targeted at this group of resistant KP. While relatively rare overall, cases of resistance emerging to the novel combination therapies (MER/VAB, CAZ/AVI, IMP/REL, AZT/AVI) during therapy have been reported (31). Resistance typically involves mutations or increased expression of a beta-lactamase (including KPC and AmpC enzymes) (32). However, loss or downregulation of OmpK36 has also been associated with increased resistance to MER/VAB (14), CAZ/AVI (33–35) and IMP/REL (34,36). Here we found that the addition of the inhibitors to the parental drugs restores susceptibility in KPC-2 producing strains with OmpK36 L3 insertions, with MICs all below resistance breakpoints. Strains not expressing OmpK36 were also susceptible to all combination drugs, with the exception of IMP/REL. Overall, these findings suggest that beta-lactamase inhibitor entry is largely OmpK36-independent. However, the MICs of CAZ/AVI, MER/VAB and AZT/AVI were modestly affected by L3 insertions, suggesting a potential contribution to increasing resistance in the presence of other mechanisms. Similarly, we found that the siderophore cephalosporin (FDC) remains effective against strains with L3 insertions, in line with its primary entry via iron uptake receptors (28).

Taken together, our data highlights OmpK36 L3 insertions as a crucial priority in the global surveillance of carbapenem resistant KP due to their high prevalence among clinical isolates, associations with large clonal expansions and relatively low fitness costs. We also propose that as genomic surveillance becomes increasingly adopted, especially in low- and middle-income countries, monitoring the diversity of such resistance mechanisms over globally-representative regions will be vital for optimisation of treatment and drug development strategies worldwide.

## Materials and Methods

### Identification and characterisation of *ompK36* among public genomes

We used a public collection of 16,086 KP genomes available in Pathogenwatch (17) (https://pathogen.watch/genomes/all?genusId=570&speciesId=573; accessed September 2021) to characterise the diversity of *ompK36* genes. The *ompK36* gene was identified in the short-read assemblies using BLASTn v2.6.0 (37) with a query gene from the reference genome, ATCC43816 (accession CP009208). To unambiguously identify *ompK36*, we required a single hit per assembly that matched ≥10% of the query length, possessed ≥90% nucleotide similarity and contained a start codon. Seaview v4.7 (38) was used to translate the nucleotide sequences to protein sequences using the standard genetic code. Non-truncated protein sequences (i.e. those with ≥95% of the query length) and the corresponding gene sequences were aligned using MUSCLE v3.8 (39). These alignments were used to identify all intact protein and gene variants present. The variants were analysed together with the metadata and genotyping data (e.g., multi-locus sequence typing and resistome data) available in Pathogenwatch. A phylogenetic tree of all intact *ompK36* gene sequences was constructed based on the variable sites using RAxML v8.2.8 (40) and visualised using Microreact v166 (41).

### Phylogenetic analysis of ST258/512, ST231 and ST16 lineages

Raw sequence reads were downloaded from the European Nucleotide Archive (ENA) for all KP genomes belonging to STs 258/512 (*n*=3673), 231 (*n*=307) and ST16 (*n*=453) in Pathogenwatch. Reads were mapped using Burrows Wheeler Aligner v0.7.17 (42) to a lineage-specific reference genome: NJST258_1 (accession CP006923) (43) for ST258/512, FDAARGOS_629 (accession NZ_CP044047) for ST231, and QS17_0029 (accession NZ_CP024038) for ST16. SNPs were identified using a pipeline comprising SAMtools mpileup v0.1.19 (44) and BCFtools v0.1.19, and pseudo-genome alignments were generated for each lineage. Individual genomes were excluded from subsequent analyses if the mean mapping coverage was <20x or if ≥25% of positions in the pseudo-genome alignment were missing data. Recombined regions were removed from the alignments and a phylogenetic tree was generated with the remaining variable positions using Gubbins v2.4.1 (45). An outgroup isolate was also included in these analyses for each lineage in order to root the phylogenetic trees (SRR5385992 from ST895 for ST258/512, ERR1216956 from ST101 for ST231, ERR1228220 from ST17 for ST16). Phylogenetic trees were visualised together with all metadata and genotyping data using Microreact v166 (41).

### Generation of OmpK36 L3 insertion mutants

Genome editing took place in the laboratory KP strain ICC8001 (a derivative of ATCC43816) using the pSEVA612S system and lambda-red fragment mediated homologous recombination as previously described. The *ompK35* gene (open reading frame) deletion was carried out in previous work (15). Mutagenesis vectors to generate the D and TD insertion were made by site directed mutagenesis using primers 1-4 (**S3 Table**). and PCR products were recircularised using KLD enzyme blend (New England Biolabs (UK)). Genome modifications were checked by sequencing of genomic *ompK36* locus PCR products generated using primers 5/6 and Sanger sequencing (Eurofins Genomics GmBH). sfGFP and mRFP1 were introduced into the silent *glmS* genomic site of strains generated in this manuscript as previously described (16).

### Purification of OmpK36 variants

OmpK36 variants were purified using our previously established protocol without any modifications (16). All OmpK36 variants were purified in 10 mM HEPES pH 7, 150 mM NaCl and 0.4% C_8_E_4_.

### Crystallisation

OmpK36_WT+TD_ and OmpK36_WT+D_ were exchanged into 10mM HEPES pH 7, 150mM LiCl, and 0.4% C_8_E_4_ prior to crystallisation. Crystals for both variants were grown from a solution containing 0.1M Lithium sulfate, 0.1M sodium citrate pH 5.6 and 12% PEG4000 at 20 °C. The crystals were cryoprotected by supplementing the crystallisation condition with 25% ethylene glycol and were frozen in liquid nitrogen. Diffraction screening and data collection were performed at Diamond Light Source synchrotron.

### Data collection and structure refinement

OmpK36_WT+D_ data to 1.78 Å were collected on I24 at Diamond Light Source at a wavelength of 0.97 Å using a Pilatus3 6M detector and processed using xia2 (46). The space group was determined to be *P1* with six copies of OmpK36_WT+D_ in the asymmetric unit. OmpK36_WT+TD_ data to 1.5 Å were collected on I03 at Diamond Light Source at a wavelength of 0.97 Å using an Eiger2 XE 16M detector and processed using xia2 (46). The resolution of both data was evaluated by half-dataset correlation coefficient in Aimless (cut-off less than 0.3) (47). The space group was determined to be *P1* with six copies of OmpK36_WT+TD_ in the asymmetric unit. Further processing was performed using the CCP4 suite (48).

Both the OmpK36_WT+TD_ and OmpK36_WT+D_ structures were determined by molecular replacement in Phaser (49) using the OmpK36_WT_ structure (PDB ID: 6RD3) (16) as a search model. Refinement of both structures was carried out in Phenix (50). After rigid body and restrained refinement electron density corresponding to the mutations and insertions were identified, built and refined. Additional density, possibly detergent or lipid molecules, that was observed on the surface of the protein was also modelled. The final OmpK36_WT+TD_ model has an R_work_ of 19% and an R_free_ of 20.5%, and the OmpK36_WT+D_ model has an R_work_ of 18.5% and an R_free_ of 21.3%, respectively.

The data collection and refinement statistics for both the OmpK36_WT+TD_ and OmpK36_WT+D_ crystals are summarised in **S2 Table**. The coordinates and structure factors of OmpK36_WT+TD_ and OmpK36_WT+D_ have been deposited to the Protein Data Bank with PDB ID codes 7PZF and 7Q3T, respectively.

### Liposome swelling assays

Liposome swelling assays were performed by reconstituting proteoliposomes with recombinant OmpK36 variants as previously described without any modifications (16).

### Antimicrobial susceptibility testing

Minimum inhibitory concentrations (MICs) were determined in triplicate by reference broth microdilution according to the ISO standard (ISO 20776-1:2019, https://www.iso.org/standard/70464.html) using sterile 96-well plates (SARSTEDT, Germany). Antibiotic and inhibitors powders were from the following sources: avibactam (AOBIOUS, U.S.A.), aztreonam (United Biotech, India), cefiderocol (Shionogi, Japan), ceftazidime, imipenem, meropenem, relebactam and vaborbactam (Merck, Germany). Cation-adjusted Mueller-Hinton broth (MHB) (Thermo Fisher Scientific, U.S.A.) was used for all agents except cefiderocol, for which iron-depleted MHB (Shionogi) was used. Inhibitors were used at fixed concentrations of 4 mg/L (avibactam, relebactam) or 8 mg/L (vaborbactam). Results were read after incubation at 35±1 °C in ambient air for 18±2 hours, and interpreted according to the EUCAST clinical breakpoints v 12.0, 2022 (https://www.eucast.org/clinical_breakpoints/), except for aztreonam/avibactam for which the EUCAST clinical breakpoint for aztreonam was used. *Escherichia coli* ATCC 25922, *Pseudomonas aeruginosa* ATCC 27853, *Klebsiella pneumoniae* ATCC 700603 and *K. pneumoniae* ATCC BAA-2814 were used as quality control strains according to EUCAST guidelines (https://www.eucast.org/ast_of_bacteria/quality_control/).

### Mouse infection

Animal work took place under the auspices of the United Kingdom Animals (Scientific Procedures) Act 1986 (License: PP7392693) and was locally approved by the institutional ethics committee. Female, BALB/c, 8-10 week old mice (Charles River, UK) were randomised into groups (co-housed in groups of 5) and acclimatised for one week before infection. Mice were housed under a 12 hour light dark cycle and had access to food and water *ad libitum*.

Inoculum was prepared from saturated overnight cultures grown in LB broth diluted in 1xPBS to a total volume of 50ul. This was delivered via intratracheal intubation (Kent Scientific) under ketamine (80mg/kg) and medetomidine (0.8mg/kg) anaesthesia. Monitored recovery from anaesthesia took place at 32-35 °C following the administration of atipamezole (1mg/kg) reversal. Inoculum size was verified (+/-10%) by enumeration on agar plates.

Bacterial enumeration at experimental end-point took place on LB agar plates supplemented with 50mcg/ml rifampicin. Blood samples were collected at end-point by cardiac puncture under terminal anaesthesia (ketamine 100mg/kg, medetomidine 1mg/kg). Lungs were excised post mortem *en bloc* followed by homogenisation. Samples were 10-fold serially diluted before plating and agar plates incubated overnight at 37 °C. In competition assays plates were UV transilluminated to determine colony fluorescence.

### Serum cytokine bead assay

Serum cytokine levels were determined using a beads-based immunoassay with a custom-designed mouse cytokine panel (LEGENDplex, BioLegend) following the manufacturer’s instructions. Cytokine measurements were acquired using a FACSCalibur flow cytometer (BD Biosciences), and flow cytometry analyses were performed using LEGENDplex data analysis software suite (https://www.biolegend.com/en-us/legendplex/software). All values were above the detection limit for both serum IL-6 and CXCL-1 (KC) levels and only 7/47 individual values (<15%; maximum of 2 per experimental group) were below the limit of detection when analysing TNF levels. These TNF values below the detection limit were assumed to be the lowest value detectable by the assay for statistical analysis and were displayed as 1 for easy visualisation in graphs (**Figure 5E**).

### Neutrophil quantification in lung homogenate by flow cytometry

1 ml of complete RPMI supplemented with penicillin and streptomycin solution (final concentration of 100 U and 100 μg/ml respectively), Liberase TM (0.13 mg/ml final, Roche) and DNaseI (10 μg/mL; Sigma-Aldrich) was added to the lung homogenate after removal of 30 μl for lung CFU calculations. The homogenate was then incubated for 40 mins on a shaker at 37 °C to allow for enzymatic tissue disruption and cell dissociation. After this incubation, the samples were placed on ice to interrupt the enzymatic digestion and the enzymes were further inhibited by adding EDTA (final concentration 10 mM; Gibco). The samples were run through a 70 μm cell strainer to obtain a uniform single-cell suspension, spun down and resuspended in 600 μl of complete RPMI. From here onwards, steps were carried out on ice to preserve cell viability.

For staining, ~5×10^6^ cells (around 120 μl) from each sample were added per well in a 96-well V-bottom plate. Dead cells were routinely excluded with Zombie Aqua Fixable Dead Cell Stain (ThermoFisher Scientific). Single cell suspensions were incubated with Fc block (Miltentyi Biotec) in FACS buffer (1% BSA, 2mM EDTA in DPBS, Sigma), followed by staining with anti-CD11b-PerCP-Cy5.5 (#45-0112-82, ThermoFisher Scientific) and anti-Ly-6G-FITC (#551460, BD Pharmingen) in FACS buffer for 30 min, all at 4°C in the dark. Fluorescence minus one (FMO) controls and a fully unstained sample were always included as controls. Stained cells were washed in FACs buffer and fixed for 20 mins at room temperature with 1% paraformaldehyde in PBS; fixed cells were kept in the dark at 4°C until analysis, usually the following day. On the day of the analysis, single stain controls for compensation were prepared using VersaComp beads (Beckman Coulter) and ArC Amine Reactive Compensation beads (ThermoFisher Scientific) and a known volume of CountBright absolute beads (ThermoFisher Scientific) was added to each sample before running the samples. Flow cytometry analysis on 50000 live cells was performed on a BD LSRFortessa cell analyzer (BD Biosciences). Data were analysed using FlowJo software (Tree Star). Neutrophils were defined as CD11b+ Ly6G+ live cells and absolute numbers of cells in the sample were calculated using the numbers of CountBright absolute beads counted following the manufacturer’s instructions.

### Data availability

All genome data used in this study is publicly available in Pathogenwatch (https://pathogen.watch/genomes/all?genusId=570&speciesId=573) and the European Nucleotide Archive (see **S1 Table** for accession numbers). Structural data corresponding to OmpK36_WT+TD_ and OmpK36_WT+D_ have been deposited in the Protein Data Bank with PDB ID codes 7PZF and 7Q3T, respectively.

## Acknowledgements

We would like to thank the Pathogen Informatics group from the Wellcome Sanger Institute for informatics support.

## Author contributions

SD and JLCW conceived the study. SD performed bioinformatics and genomic analyses. JLCW performed the molecular biology and biochemistry experiments. JLCW, JSG and WWL performed the animal experiments under the supervision of GF. JSG performed cytokine measurements. H-SK and KB performed the crystallography and liposome swelling assays. FM, TG and GMR performed the meropenem resistance assays. Data analysis was carried out by JLCW, SD, H-SK, KB and JSG. JLCW, SD and GF wrote the manuscript. All authors reviewed and edited the manuscript.

